# Novel derivatives of BCV and (S)-HPMPA inhibit orthopoxviruses and human adenoviruses more potently than BCV

**DOI:** 10.1101/2024.08.30.610570

**Authors:** Yifan Zhang, Yanmin Wan, Cuiyuan Guo, Zhaoqin Zhu, Chao Qiu, Jiasheng Lu, Yanan Zhou, Jiaojiao Zheng, Fahui Dai, Xiaoyang Cheng, Kunlu Deng, Wanhai Wang, Youchun Wang, Wenhong Zhang

**Author notes:** These authors contribute equally: Yifan Zhang, Yanmin Wan, Cuiyuan Guo, Zhaoqin Zhu and Chao Qiu. These authors jointly supervised this work: Yanmin Wan,; Youchun Wang,; Wenhong Zhang.

## Abstract

BCV and tecovirimat are the only two chemical drugs that have been approved to treat smallpox and can be requested for Mpox treatment through a single-patient Emergency Investigational New Drug (EIND) application. Disappointedly, the efficacy of tecovirimat manifested in a recent clinical trial is far from being satisfactory, while the clinical efficacy of BCV is still inconclusive. Given that MPXV, variola and other emerging orthopoxviruses are posing serious threats to global health, it is urgent to develop better therapeutics. In this study, we tested the antiviral effects of three novel prodrugs, which were designed based on previously reported parent drugs, either (S)-HPMPC (cidofovir) or (S)-HPMPA. We found that one of the (S)-HPMPA-based prodrugs, ODE-(S)-HPMPA formate, exhibited significantly better anti-orthopoxvirus activity than BCV both in vitro and in vivo, which also inhibited human adenovirus type 2 and type 21 more efficiently than BCV. Most strikingly, the EC_50_ and EC_90_ of ODE-(S)-HPMPA formate against MPXV were more than 40-fold lower than those of BCV. In contrast, we observed that the anti-HSV-1 activities of the (S)-HPMPA-based prodrugs were less effective than those of the cidofovir-based prodrugs (BCV and BCV formate), especially in vivo. Moreover, we showed for the first time that cytidine and adenine analog combined therapies could provide mice with complete protection against lethal challenges of both vaccinia and HSV-1. Collectively, we propose that both the ODE-(S)-HPMPA formate and the BCV/ODE-(S)-HPMPA formate combination are worth further investigations for their potential clinical applications.

## Introduction

Viral polymerases are the most sought-after targets for developing anti-viral drugs [1–3]. Few dozens of viral RNA or DNA dependent polymerase inhibitors have been licensed for the treatment of human viral diseases, including brincidofovir (BCV, a prodrug of cidofovir), which is the second FDA approved drug for the treatment of smallpox infection [3, 4]. Given that variola and monkeypox virus (MPXV) are phylogenetically similar, the U.S. CDC suggests that BCV can be requested for the treatment of human Mpox disease in adults and pediatric patients through a single-patient Emergency Investigational New Drug (EIND) application.

BCV is a lipid conjugate of the nucleotide analogue cidofovir with broad-spectrum anti-double stranded DNA virus activity [5]. Data of previous studies suggested that the in vitro 50% inhibitory concentration (IC_50_) of BCV against MPXV was relatively higher than that of tecovirimat [6–8], a specific inhibitor of orthopoxviruses targeting viral p37 protein orthologs [9]. Both compounds have been used to treat human Mpox disease [8, 10], but BCV can elevate the transaminases in the liver [11–13]. Tecovirimat has a relatively better safety profile [14] and is currently recommended by the U.S. CDC to be the first line treatment for MPXV infections in people. But at the same time, the U.S. CDC also alerts in the Guidance for Tecovirimat Use (https://www.cdc.gov/poxvirus/mpox/clinicians/Tecovirimat.html) that tecovirimat has a low barrier to viral resistance. A single amino acid mutation to the orthopoxviral p37 protein can lead to resistance to tecovirimat [15], which may be especially prong to occurrence among severely immunocompromised Mpox patients under multiple courses of tecovirimat treatment [16, 17]. Besides, according to data released from a clinical trial in the Democratic Republic of the Congo, tecovirimat neither accelerated recovery nor reduced mortality rate compared to placebo [18].

Mutations associated with cidofovir resistance have also been identified in multiple orthopoxviruses [8, 19] and human cytomegalovirus [20]. And mutations potentially associated with CDV resistance among the MPXVs of 2022 outbreak have been reported, too [8]. The prevalence of mutated MPXV may further rise once BCV was more widely and frequently used, which is also true for other orthopoxviruses because the mechanism of CDV resistance seems to be shared among the genus [21]. Given the pacing threats posed by MPXV, variola and other emerging orthopoxviruses to human health [22–25], it is urgent to develop safer and more effective drugs.

CDV [(S)-1-(3-hydroxy-2-phosphonylmethoxypropyl)cytosine, HPMPC] and its prodrug BCV exemplify that the (S)-HPMP nucleoside can be a promising class of broad spectrum anti-viral compounds [5]. In fact, many others candidate molecules in this class have been synthesized and tested for anti-viral effects. (S)-HPMPA [(S)-9-(3-hydroxy-2-phosphonomethoxypropyl)-adenine] is one of these candidates, which showed broad spectrum anti-viral activities in earlier studies [5, 26]. However, because of the poor oral bioavailability, (S)-HPMPA is considered as not suitable for clinical use. Great efforts have been made to modify (S)-HPMPA in order to increase their oral bioavailabilities and anti-viral activities [27–35]. But only few candidate compounds, such as ODE-(S)-HPMPA and HDP-(S)-HPMPA, displayed promising in vivo anti-orthopoxvirus effects [30, 33]. In this work, we synthesized two novel (S)-HPMPA prodrugs, ODE-(S)-HPMPA formate and HDP-(S)-HPMPA formate, and a new CDV prodrug, BCV formate, and tested their anti-viral effects in parallel with BCV both in vitro and in vivo.

## Materials and methods

### Ethical Statement

Experiments using mice were approved by the Research Ethics Review Committee of the Shanghai Public Health Clinical Center Affiliated to Fudan University. Experiments using live vaccinia, HSV-1 and adenoviruses were conducted in a BSL-2 or ABSL-2 lab. Measurements of in vitro inhibition of MPXV were performed in a BSL-3 lab.

### Antiviral compounds

BCV (CAS number 444805-28-1, purity 99.82%) was purchased from Shanghai Haohong Scientific Co., Ltd.. Three candidate compounds were synthesized by our collaborator, Risen (Shanghai) Pharma Tech Co., Ltd. Compound BCV formate (purity 97.86%) is a derivative of cidofovir. Compound ODE-(S)-HPMPA formate (purity 95.03%) and compound HDP-(S)-HPMPA formate (purity 96.97%) are prodrugs of (S)-HPMPA. The molecular formulas of these compounds and their active metabolites are shown in Figure 1.

**Figure 1.**
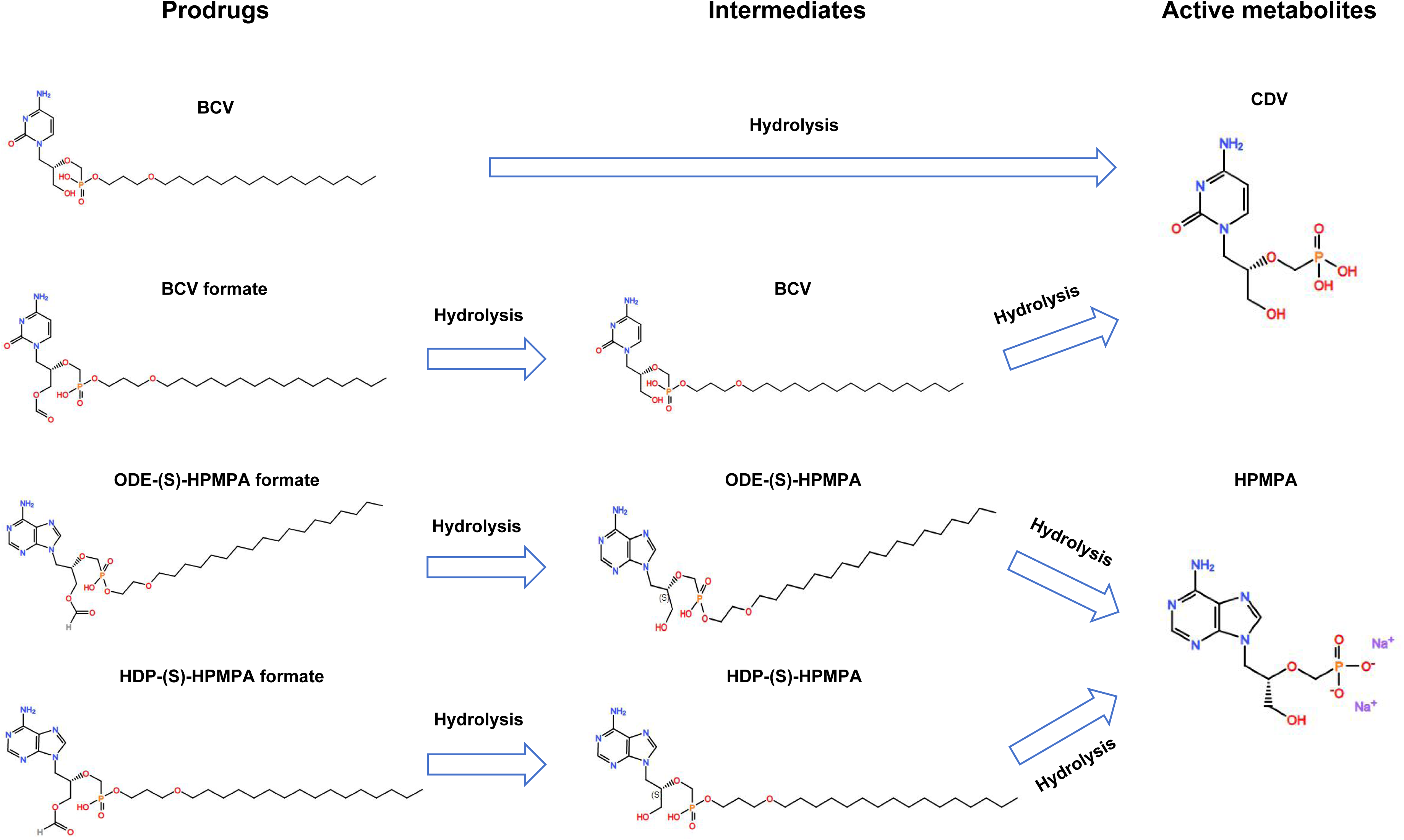
Illustration of the molecular formulae and the theoretical metabolic pathways of BCV and the newly synthesized prodrugs. After two steps of hydrolysis, BCV formate can be converted to the same active parent drug (CDV) with BCV. ODE-(S)-HPMPA formate and HDP-(S)-HPMPA formate also take two steps of hydrolysis to turn into HPMPA.

### Virus propagation and titration

Vaccinia Tiantan strain was propagated in Vero cells. Briefly, confluent Vero cells plated in 100mm×10mm petri dishes were inoculated with Tiantan vaccinia at a multiplicity of infection (MOI) of 0.01. After 2 hours absorption at 37°C with 5% CO_2_, the inoculum was removed and 10ml of fresh maintenance medium (DMEM containing 3% FBS and 1% PS) was added to each dish. The cell cultures were maintained at 37°C with 5% CO_2_ for another 2 days. After that, propagated vaccinia viruses were harvested by three rapid freeze-thaw cycles of the infected cells. HSV-1 propagation followed a similar protocol except that the inoculum was adsorbed for 1 hour with gentle agitation at room temperature and the propagated virus was harvested from the culture supernatant by centrifuging at 34,000g for 2 hours at 4°C [36]. MPXV was sourced from National Kunming High-level Biosafety Primate Research Center, Institute of Medical Biology, Chinese Academy of Medical Sciences and Peking Union Medical College. Viruses were amplified in Vero cells, cultured in DMEM supplemented with 2% FBS and 1% PS. Harvested vaccinia, MPXV and HSV-1 viruses were titrated using plaque assays. Briefly, confluent monolayers of Vero cells in 24-well plates were infected with 10-fold serial dilutions of each virus, ranging from 10^−1^ to 10^−9^. For vaccinia viruses and MPXV, the plates were incubated at 37°C with 5% CO_2_ for 2 hours. For HSV-1 (strain-17), the plates were gently agitated at room temperature for 1 hour. After absorption, the cells were overlaid with 800 μL of overlay medium (DMEM containing 1% FBS and 1.25% methyl cellulose) and incubated at 37°C with 5% CO_2_ for 4 days. Post-incubation, the plates were fixed with 4% neutral formalin and stained with crystal violet. Plaques were visually counted, and the viral titers were calculated and expressed as plaque-forming units (PFU) per milliliter.

HAdV-B21 and C2 were isolated from nasopharyngeal swab specimens of HAdV-positive patients and propagated in HEp-2 cells. Briefly, HEp-2 cells were plated in 100mm×10mm petri dishes and inoculated with hAdVs. After a 1-hour adsorption at 37° C with 5% CO_2_, the inoculum was removed, and 10 mL of maintenance medium was added to each dish. The cultures were maintained at 37°C with 5% CO_2_ until more than 80% of the cells exhibited cytopathic effects (CPE). Propagated hAdVs were harvested by three rapid freeze-thaw cycles of the infected cells.

The stocks of hAdVs were titrated using the TCID_50_ assay. Briefly, confluent monolayers of Vero cells in 96-well cell culture plates were infected with 10-fold serial dilutions of the hAdVs. The wells were monitored for the presence of CPE and the infectious adenovirus titer was determined as TCID_50_ per mL using the Spearman-Karber method.

### Cytotoxicity assays

The cytotoxicities of candidate compounds were determined using a Promega CellTiter-Glo® Luminescent Cell Viability Assay kit (Cat#G7571, Madison, WI, USA) according to the manufacturer’s instructions. Briefly, Vero cells were seeded in 96-well plates and incubated overnight. Then, cells were exposed to either the solvent control (DCM:MeOH, v:v=1:3) or diluted candidate compounds. Each concentration was tested in quadruplicate. The plates were incubated for 72 hours at 37°C with 5% CO_2_. Subsequently, reconstituted Steady-Glo reagent was added and assay plates were incubated for 10 min at room temperature. The luminescence was measured using a luminescence microplate reader (GloMax® Navigator Microplate Luminometer, Promega, USA). Wells containing culture medium without cells served as blank controls. Cell viabilities were calculated using the following equation: Cell viability (%) = (drug treated well - blank) / (solvent treated well - blank) × 100%.

### In vitro viral inhibition assays

Twenty PFU of orthopoxvirus (Tiantan vaccinia or MPXV) or HSV-1 virus suspended in 200 μL maintenance medium were transferred onto confluent Vero cell monolayers in 24-well plates. For vaccinia virus, the plates were incubated at 37°C with 5% CO_2_ for 2 hours. For HSV-1, the plates were gently agitated at room temperature for 1 hour. 800 μL of overlay-medium containing DCM:MeOH (v:v=1:3) or serially diluted candidate compounds was added to each well after removing the inoculums. The plates were then incubated for 4 days at 37°C with 5% CO_2_. Finally, the plates were fixed with 4% neutral formalin and stained with crystal violet. The number of plaques in each well was visually counted. All compounds were tested in quadruplicates at each dilution. The viral inhibition rate was calculated using the formula: Inhibition rate (%) = (1 − the average plaque number of the sample wells / the average plaque number of the solvent wells) × 100%.

In vitro inhibition of hAdV B21 and C2 were performed as described previously [37]. Briefly, 25,000 Vero cells were plated in each well of a 96-well cell culture plate, followed by addition of diluted adenovirus (with 50 and 200 TCID_50_/well for B21 and C2, respectively) and serially diluted candidate compounds. The solvent was used as the non-treated control. The plates were then incubated at 37°C with 5% CO_2_ for 4 days. Cytopathology was quantitively assessed using a Promega CellTiter-Glo® Luminescent Cell Viability Assay kit. The viral inhibition rate was calculated using the formula: inhibition rate (%) = (drug treated well – blank) - (non-treated control well – blank) × 100%.

### In vivo evaluation of anti-viral effects

We established mouse infection models to evaluate the anti-viral effects of the candidate compounds against Tiantan vaccinia and HSV-1, respectively. Vaccinia Tiantan strain (6×10^6^ PFU) was suspended in 40 μL 1×PBS and instilled intranasally into 4-week-old male BALB/c mice under transient anesthesia of inhaled isoflurane (Cat# R510-22, Ryward RWD). For HSV-1 infections, 4-week-old female BALB/c mice were infected intraperitoneally with 1×10^8^ PFU of HSV-1 suspended in 200 μL of 1×PBS. After infection, mice were weighed daily to monitor disease progression and mice that lost ≥30% of their initial body weight were deemed ethically dead. All mice were daily monitored to the end of the experiment and biological death cause by vaccinia infection were also recorded. For treatment experiments, mice were given equal moles of candidate compounds dissolved in saline containing 5% DMSO, 10% Solutol, and 17% SBE-β-CD via oral gavage at days 1, 3 and 5 post-infection as specified in Figure 4-6. Control animals received an equivalent volume of the vehicle via the same gavage method. Infected mice were euthanized at timepoints indicated in Figure 4-6. Samples of multiple organs were collected for viral titration, host response assay or pharmaceutical analysis.

### Pharmaceutical analyses of mouse tissue samples

Anticoagulated mouse blood samples were collected and centrifuged at 3200 rpm for 10 minutes at 4°C to separate the plasma, which was then aliquoted and stored at −80°C until analysis. Tissue samples, including spleen, brain, liver, kidney, lung, and testes, were homogenized and centrifuged at 4100g for 15 minutes to obtain supernatants for pharmacokinetic analysis. The concentrations of the candidate compounds, intermediate metabolites, and active metabolites in both plasma and tissue samples were determined using a validated liquid chromatography-tandem mass spectrometry (LC-MS/MS) method, as described in the literature [38]. The lower limits of quantitation (LLOQ) were 1 ng/mL in plasma, and in tissues, the LLOQs were 10 ng/g for ODE-(S)-HPMPA and BCV, 20 ng/g for CDV, and 50 ng/g for HPMPA and HDP-(S)-HPMPA.

### Titration of Tiantan vaccinia and HSV-1 in mouse organs

Tissue samples were pre-cooled on ice and homogenized using high-throughput tissue grinding machine (Cat# Scientz-192, NingBo Scientz Biotechnology Co., Ningbo City, China). After that, the homogenates were centrifuged at 2000g for 10 minutes and the supernatants were collected for viral titration following similar procedures to those described above for in vitro viral titration. Briefly, confluent monolayers of Vero cells in 24-well plates were inoculated with 10-fold serially diluted supernatants of tissue homogenates. After incubation, the wells were overlaid with 800 μL of overlay medium and incubated for 4 days. Finally, the plates were fixed with neutral formalin and stained with crystal violet, and plaques were counted to determine viral titers.

### Histological examination

Tissue samples of multiple mouse organs were collected at 7 days post Tiantan vaccinia infection. Collected tissues were fixed in 4% neutral paraformaldehyde for 2 hours, followed by standard processing and embedding in paraffin. Tissue sections were stained with hematoxylin and eosin (H&E). The stained slides were analyzed using the KF-PRO-120 digital pathology slide scanner (KFBIO) to assess tissue morphology and the extent of pathological changes. The images were analyzed and exported by K-VIEWER1.5.5.6 software.

### Detection of cytokine transcription in mouse lung

Total RNA was extracted from mouse lung tissues collected at multiple timepoints post Tiantan vaccinia infection using Trizol reagent (Cat# 15596026CN, Life Technologies). 1 μg of total RNA from each sample was reversely transcribed using the HiScript II Q RT SuperMix for qPCR (+gDNA wiper) (R223-01, Vazyme, China). Real-time PCR was conducted with the TB Green® Premix Ex Taq™ II (Tli RNase H Plus) (Cat# RR820Q, TaKaRa) using an ABI 7500 real-time fluorescence quantitative PCR (qPCR) instrument (Thermo Fisher Technology Co., LTD., Shanghai, China). Transcription levels of cytokine genes relative to the GAPDH gene were calculated using the 2^−ΔCt^ method, where ΔCt = Ct of the target gene - Ct of GAPDH. The primers of mouse cytokines and GAPDH genes used for qRT-PCR are listed in Supplemental table 1, which were synthesized according to a previous work [39].

### Determination of Alanine Aminotransferase (ALT) in murine serum

Peripheral blood was collected from mice at 2 days post the third dose of drug treatment. Serum was separated after clotting at room temperature for 1 hour by centrifuging at 1000g for 10 minutes at 4°C. Serum ALT activity was determined using a commercialized ELISA kit following the manufacturer’s protocol (Cat# JL12668, Jianglai Biotechnology, Shanghai, China). Optical density was measured at 450 nm using a microplate reader (Cat# 800TS, Biotek, USA).

### Statistical methods

Statistical analyses were conducted using GraphPad Prism 9 (GraphPad Software, USA). The normality of the data was checked before all downstream statistical analyses. Comparisons between two groups were performed by the method of t-test. Differences among multiple groups were compared by the method of one-way ANOVA. EC_50_ and EC_90_ values were calculated using a four-parameter variable slope non-linear regression method. Survival curves were compared by the Cox-Mantel test. P ≤ 0.05 was considered as statistically significant.

## Results

### In vitro cytotoxicity and in vivo pharmacokinetics of candidate compounds

Synthesized compounds (BCV formate, ODE-(S)-HPMPA formate and HDP-(S)- HPMPA formate) were dissolved in DCM:MeOH (v:v=1:3) at a maximum concentration of 10μM. Cytotoxicity assays showed that all compounds were well tolerated at concentrations of 5μM and 10μM in Vero cells (Figure 2A and 2B).

**Figure 2.**
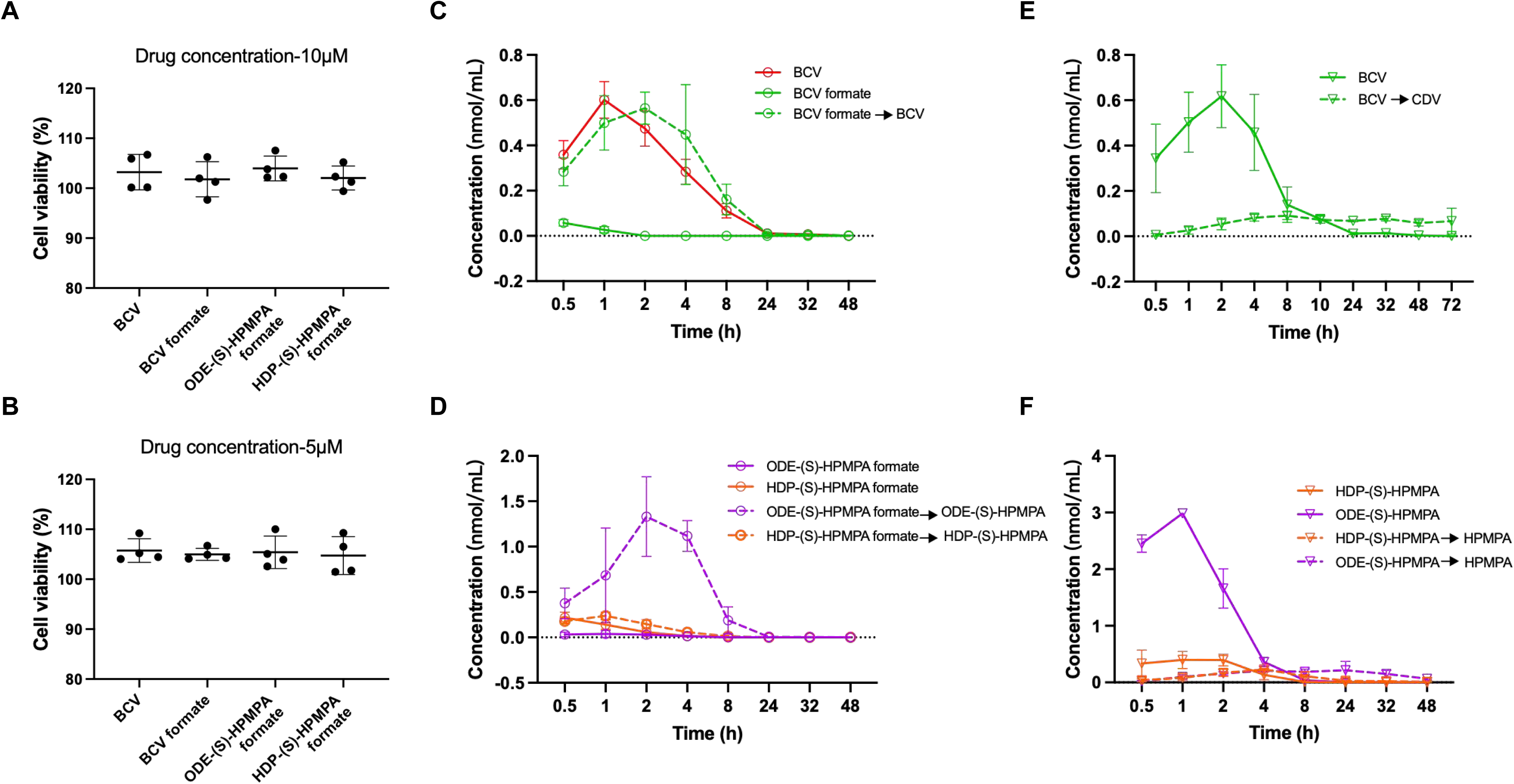
In vitro cytotoxicity and in vivo pharmacokinetic analyses of BCV and the candidate compounds. **(A-B**) Comparisons of cytotoxicities between candidate compounds and BCV. Limited by the solubility of BCV formate, ODE-(S)-HPMPA formate and HDP-(S)-HPMPA formate, the highest concentration employed in the cytotoxicity assay was 10μM. All compounds were tested in quadruplicated wells and the solvent was used as the negative control to calculate the percentage of cell viabilities. Data are presented as mean ± SD. (**C-D**) Plasma concentrations of BCV, BCV formate, ODE-(S)-HPMPA formate and HDP-(S)-HPMPA formate, as well as their respective one-step hydrolytic metabolites CDV, BCV, ODE-(S)-HPMPA and HDP-(S)-HPMPA, were measured at various time points following a single oral gavage dose (3 mice per group). (**E-F**) Plasma concentrations of BCV, ODE-(S)-HPMPA, and HDP-(S)-HPMPA, as well as their hydrolytic metabolites (CDV and HPMPA) were measured at various time points following a single oral gavage dose (3 mice per group).

To facilitate in vivo antiviral activity evaluation, we monitored the kinetics of absorption and metabolism for each compound after a single dose of oral gavage in mice. Our data showed that all compounds or their metabolites were detectable at a half hour post oral administration (Figure 2C and 2D), suggesting that they could be quickly assimilated into blood. After absorption, BCV formate, ODE-(S)-HPMPA formate and HDP-(S)- HPMPA formate were instantly hydrolyzed into BCV, ODE-(S)-HPMPA and HDP-(S)- HPMPA, respectively. And the concentrations of these metabolites in blood peaked around 1 to 2 hours post oral administration (Figure 2C and 2D). Then, BCV was gradually hydrolyzed into CDV within 24 hours post oral administration (Figure 2E). While ODE-(S)-HPMPA and HDP-(S)-HPMPA were hydrolyzed faster into HPMPA, which lasted only 8 hours (Figure 2F).

### ODE-(S)-HPMPA formate and HDP-(S)-HPMPA formate showed superior anti-orthopoxvirus activities than BCV and BCV formate

To compare the anti-orthopoxvirus effects of the candidate drugs with BCV, we firstly tested their antiviral activities against vaccinia Tiantan strain and MPXV in vitro. Our data showed that the EC_50_ and EC_90_ of BCV formate against Tiantan vaccinia were similar with those of BCV, while the EC_50_ and EC_90_ of ODE-(S)-HPMPA formate and HDP-(S)-HPMPA formate were conspicuously lower (Figure 3A). Notably, ODE-(S)- HPMPA formate displayed the lowest EC_50_ and EC_90_, which were around 15-fold lower than BCV (Figure 3A). The anti-MPXV activity of BCV was similar with its effect against Tiantan vaccinia, while the other three compounds inhibited MPXV even more potently than Tiantan vaccinia (Figure 3B). ODE-(S)-HPMPA formate demonstrated the most potent anti-MPXV effect, whose EC_50_ and EC_90_ were more than 40-fold lower than those of BCV (Figure 3B).

**Figure 3.**
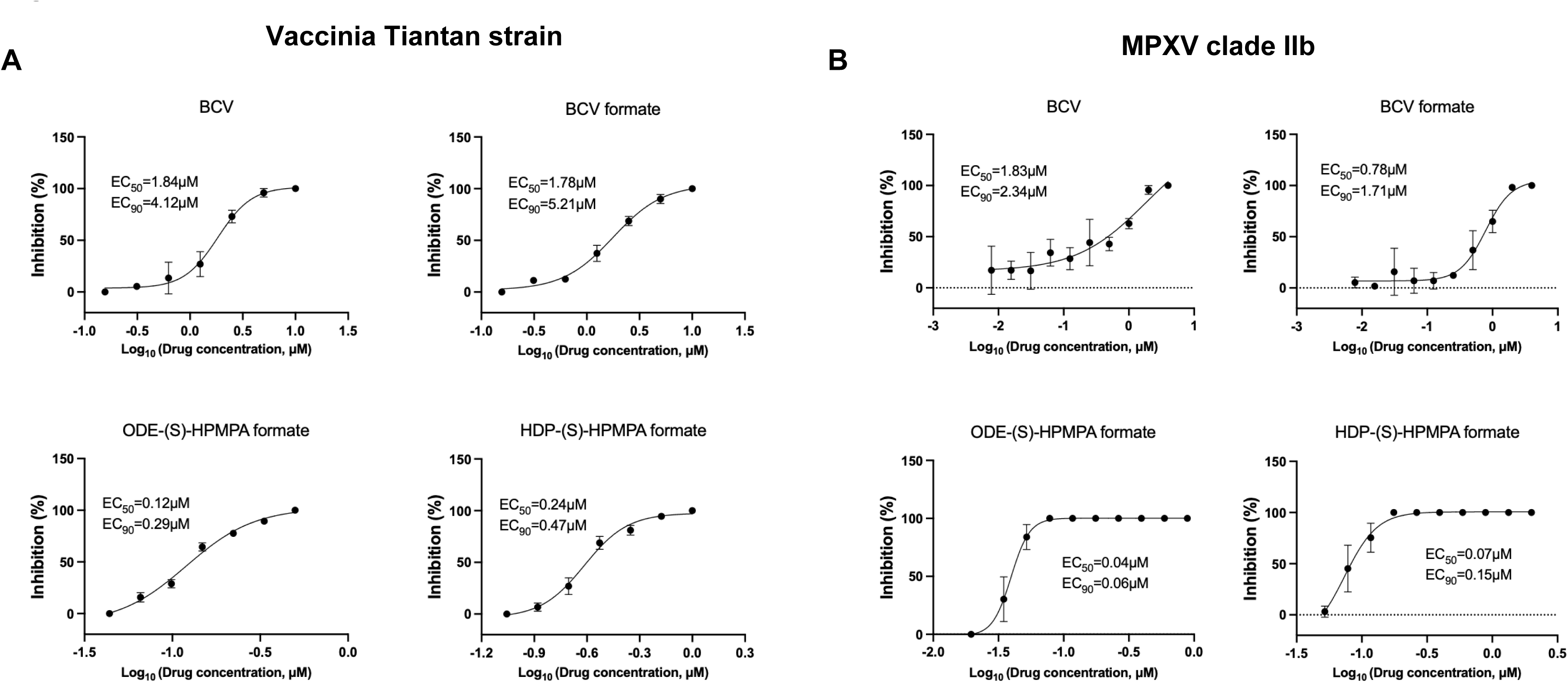
In vitro evaluations of anti-vaccinia and anti-MPXV activities of BCV and the candidate compounds. Vero cells were infected with vaccinia Tiantan strain (**A**) or MPXV (**B**) and treated with the indicated concentrations of BCV or the candidate prodrugs. All compounds were tested in triplicated wells at each concentration. The solvent was used as the non-treated control. The antiviral effects were determined by calculating the reduction in plaque formation. Data are presented as mean ± SD.

Next, we evaluated the in vivo anti-orthopoxvirus activities of the synthesized compounds using a lethal mouse model of vaccinia virus infection. As shown in Figure 4A, 4 weeks old BALB/c mice were intra nasally infected with 6×10^6^ pfu of vaccinia Tiantan strain. Equal moles of candidate drugs were given at 1 day, 3 days and 5 days post infection via oral gavage. All groups of infected mice experienced obvious body weight loss, which peaked around 7 to 8 days post infection for mice treated with BCV formate, ODE-(S)-HPMPA formate or HDP-(S)-HPMPA formate (Figure 4B). In contrast, mice received the vehicle or BCV underwent more severe weight loss and all mice in these two groups ethically died by day 8 (Figure 4B). For animals in the group treated with ODE-(S)-HPMPA formate, more than 90% of mice (10 out of 11) survived, a significant improvement in the survival rate compared with animals in all other groups (vehicle, BCV or BCV formate groups; Figure 4C). The second-best drug was HDP-(S)- HPMPA formate, which protected significantly more mice from death than vehicle and BCV (Figure 4C). BCV formate also displayed better protection, but its efficacy did not significantly differ from those of the vehicle and BCV (Figure 4C).

**Figure 4.**
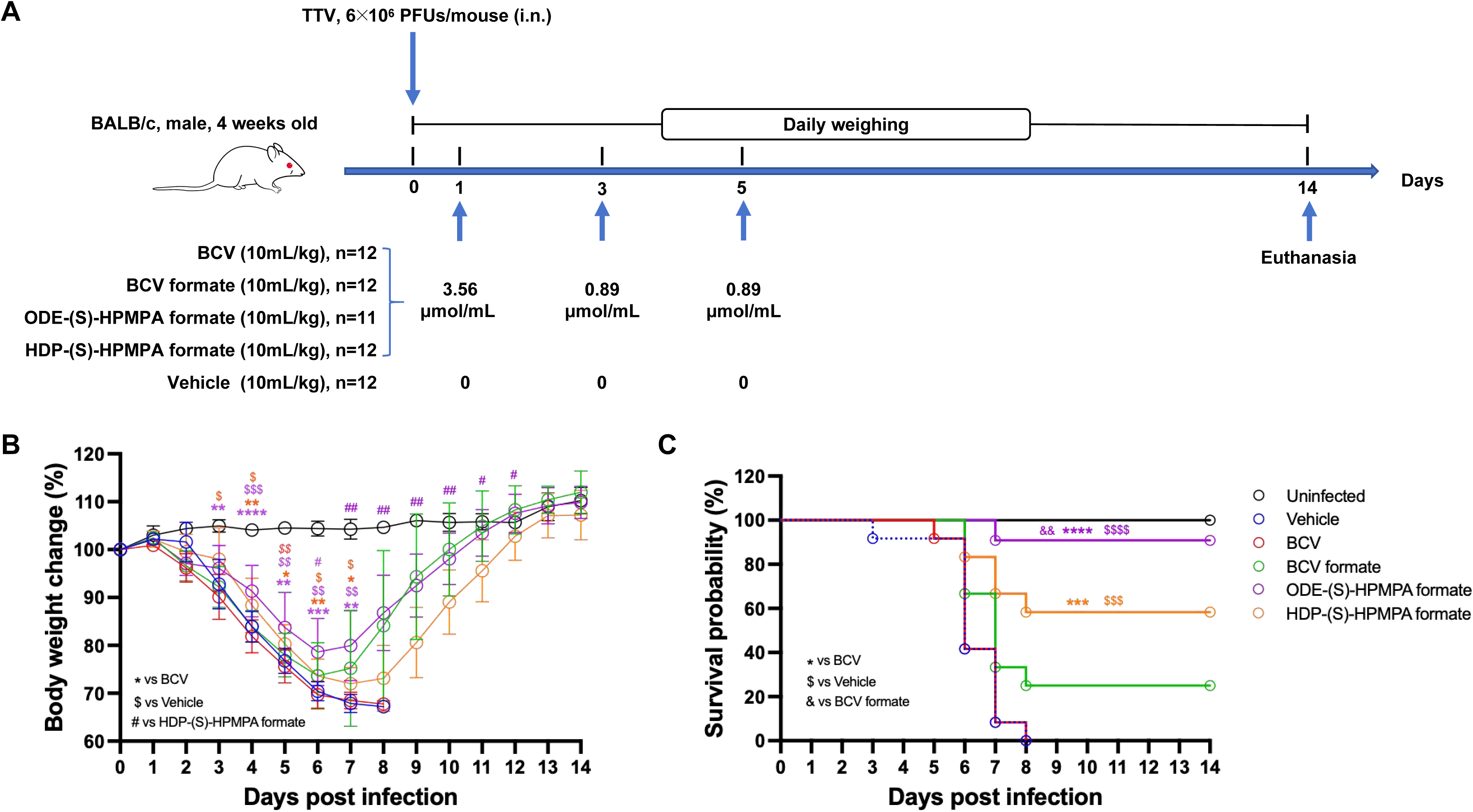
ODE-(S)-HPMPA formate and HDP-(S)-HPMPA formate showed superior therapeutic efficacies against lethal Tiantan vaccinia infection in mice. (**A**) The schematic illustration of experimental design. (**B-C**) Weight loss and survival rates of mice after being intra nasally infected with the vaccinia Tiantan strain. Mice received the solvent served as the non-treated control. Weight loss ≥30% of initial body weight was recorded as ethical death and was therefore not represented in the figure. The data of weight loss are presented as mean ± SD. Mice that lost ≥30% of their initial body weight were deemed ethically dead. Statistical significance is indicated as follows: *, $, #: p < 0.05; **, $$, ##: p < 0.01; ***, $$$, ###: p < 0.001; ****, $$$$, ####: p < 0.0001.

To characterize the in vivo anti-vaccinia virus activities further, we repeated vaccinia virus infection and drug treatment in mice, but this time 3-5 mice of each group were euthanized at indicated time points for detections of viral replication, host responses and drug metabolites (Figure 5A). We found that vaccinia virus resided mainly in lungs after intra nasal infection and significant suppression of the virus by candidate drugs could be observed in all detected tissues except kidney (Figure 5B-5G), where live vaccinia viruses were rarely detected even in the non-treated group. Compared to BCV, BCV formate, ODE-(S)-HPMPA formate and HDP-(S)-HPMPA formate tended to decrease vaccinia virus titers in lungs, but statistical significance was only observed between BCV and ODE-(S)-HPMPA formate at day 3 and between BCV and HDP-(S)- HPMPA formate at day 5 (Figure 5B). Histopathology examination showed that ODE- (S)-HPMPA formate and HDP-(S)-HPMPA formate diminished the size of lung lesions caused by vaccinia infection compared to BCV and BCV formate at 7 days post infection (Figure 5H). Meanwhile, no signs of infection-caused or drug-related pathology were observed in other organs, including the kidney, spleen, testis, liver and brain (Supplementary figure 1). We also monitored the cytokine responses in lungs at 3, 5, 7, 10 and 14 days post infection. We found that different drugs impacted lung inflammation differentially (Supplementary figure 2). Of note, at 7 days post vaccinia infection (approximately the nadir of mouse body weight change), the transcription levels of IL-6 in lungs of mice treated with ODE-(S)-HPMPA formate were significantly lower than those of mice treated with vehicle or BCV (Supplementary figure 2A). While, the levels of IL-12 and TNF-a transcripts in lungs of mice treated with ODE-(S)-HPMPA formate or BCV formate were significantly higher than those of mice treated with vehicle or BCV (Supplementary figure 2B and 2C). These findings were partially in accordance with a previous study suggesting that HPMPA could activate TNF-a secretion [40].

**Figure 5.**
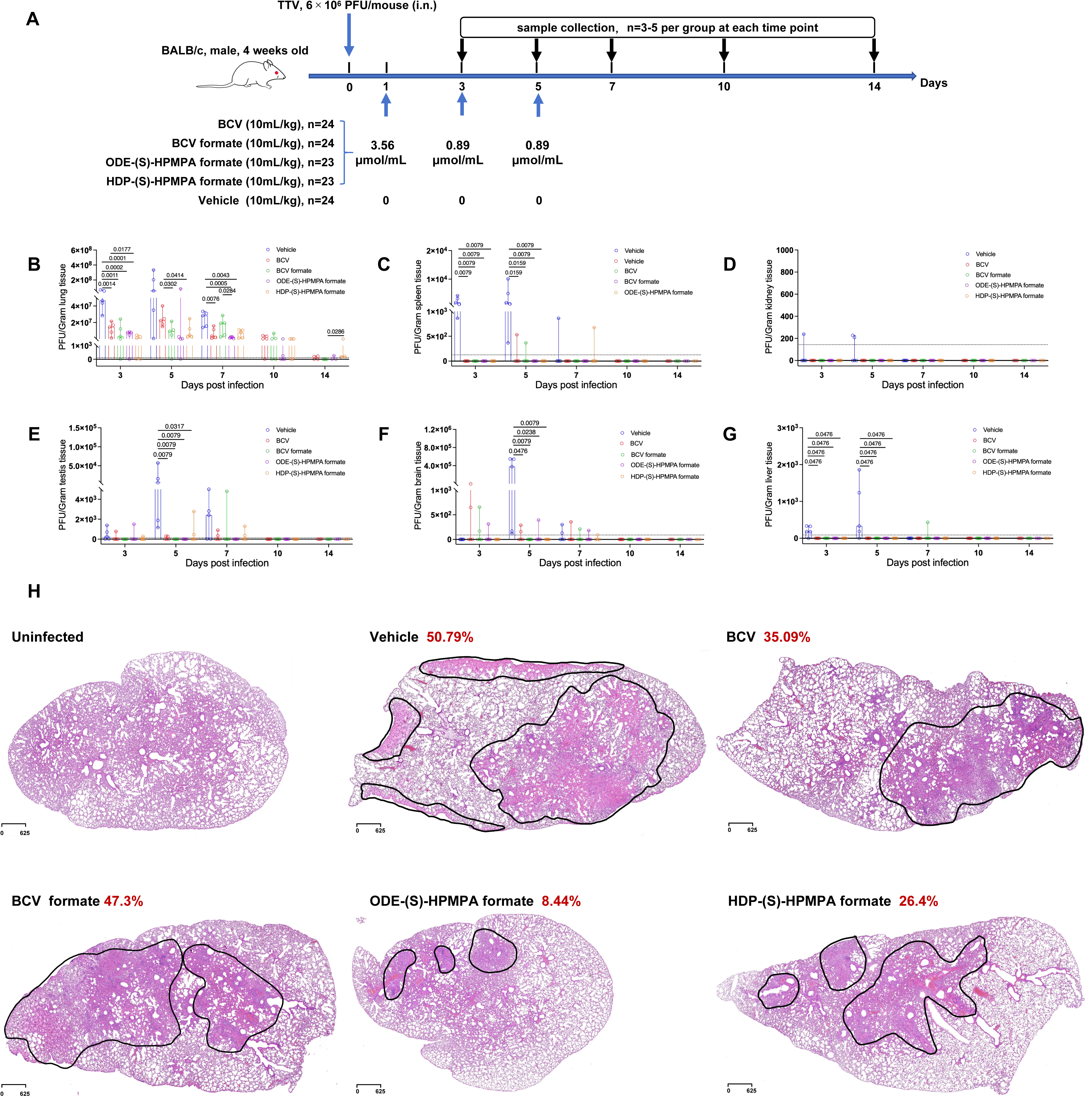
ODE-(S)-HPMPA formate and HDP-(S)-HPMPA formate more effectively reduced lung vaccinia virus titers and mitigated pathological severity compared to BCV and BCV formate. (**A**) The schematic illustration of experimental design. (**B-G**) Viral titers in the lung, spleen, kidney, brain, testis, and liver were measured by plaque assays at 3, 5, 7, 10, and 14 days post-infection, respectively. (**H**) Histopathological examination of lung tissue at 7 days post infection. Black outlines indicate areas of observed pathology, and the percentages of the lesion area relative to the total area of the tissue are denoted in red.

We noticed that livers contained the highest concentrations of drug metabolites during the whole process of drug treatments (Supplementary figure 3). Hence, we assessed the liver toxicities of candidate drugs via measuring the serum ALT levels at day 7 (2 days post the third dose of drug treatment). The results showed that the average levels of plasma ALT ranked as follows: HDP-(S)-HPMPA formate > BCV formate > BCV > vehicle > ODE-(S)-HPMPA formate > uninfected control (Supplementary figure 4). Of note, the ALT levels of BCV, BCV formate and HDP-(S)-HPMPA formate treated mice were significantly higher than that in uninfected controls, while no significant difference was observed among mice of ODE-(S)-HPMPA formate treated group, vehicle group, and control group (Supplementary figure 4).

### ODE-(S)-HPMPA formate and HDP-(S)-HPMPA formate could suppress HSV-1 both in vivo and in vitro, but were less effective than BCV and BCV formate in protecting mice against lethal infection

As NAs usually have broad spectrum antiviral activity [41], we next compared the anti-HSV-1 activities between the candidate drugs and BCV. The results of in vitro antiviral assays indicated that the EC_50_ and EC_90_ of BCV formate and ODE-(S)-HPMPA formate against HSV-1 were close to those of BCV, while the EC_50_ and EC_90_ of HDP-(S)- HPMPA formate were slightly higher (Figure 6A). To validate this observation, we tested their efficacies against HSV-1 lethal infection in vivo. As shown in Figure 6B, BALB/c mice were intra peritoneally infected with 1×10^8^ pfu of HSV-1 (strain-17) followed by the same regimen of drug treatment with vaccinia infected mice. Our data showed that BCV and BCV formate alleviated mouse body weight loss, whereas ODE-(S)-HPMPA formate and HDP-(S)-HPMPA formate apparently did not (Figure 6C). Albeit ODE-(S)- HPMPA formate and HDP-(S)-HPMPA formate provided significant protection against lethal HSV-1 infection than the vehicle, their protection rates were significantly lower than those of BCV and BCV formate (Figure 6D). It is intriguing to see that the adenine analogs (ODE-(S)-HPMPA formate and HDP-(S)-HPMPA formate) were more effective in protecting mice against vaccinia infection while less effective towards HSV-1 infection compared to cytidine analogs (BCV and BCV formate). We further compared the genomic DNA base composition of orthopoxvirus with that of HSV and found that orthopoxviruses contain obviously less GC than HSV (Supplementary figure 5 and Supplementary table 2). This difference might lead to that adenine analogs would have more chance to be incorporated into orthopoxvirus genome, while the cytidine analogs would be more frequently incorporated into HSV genome.

**Figure 6.**
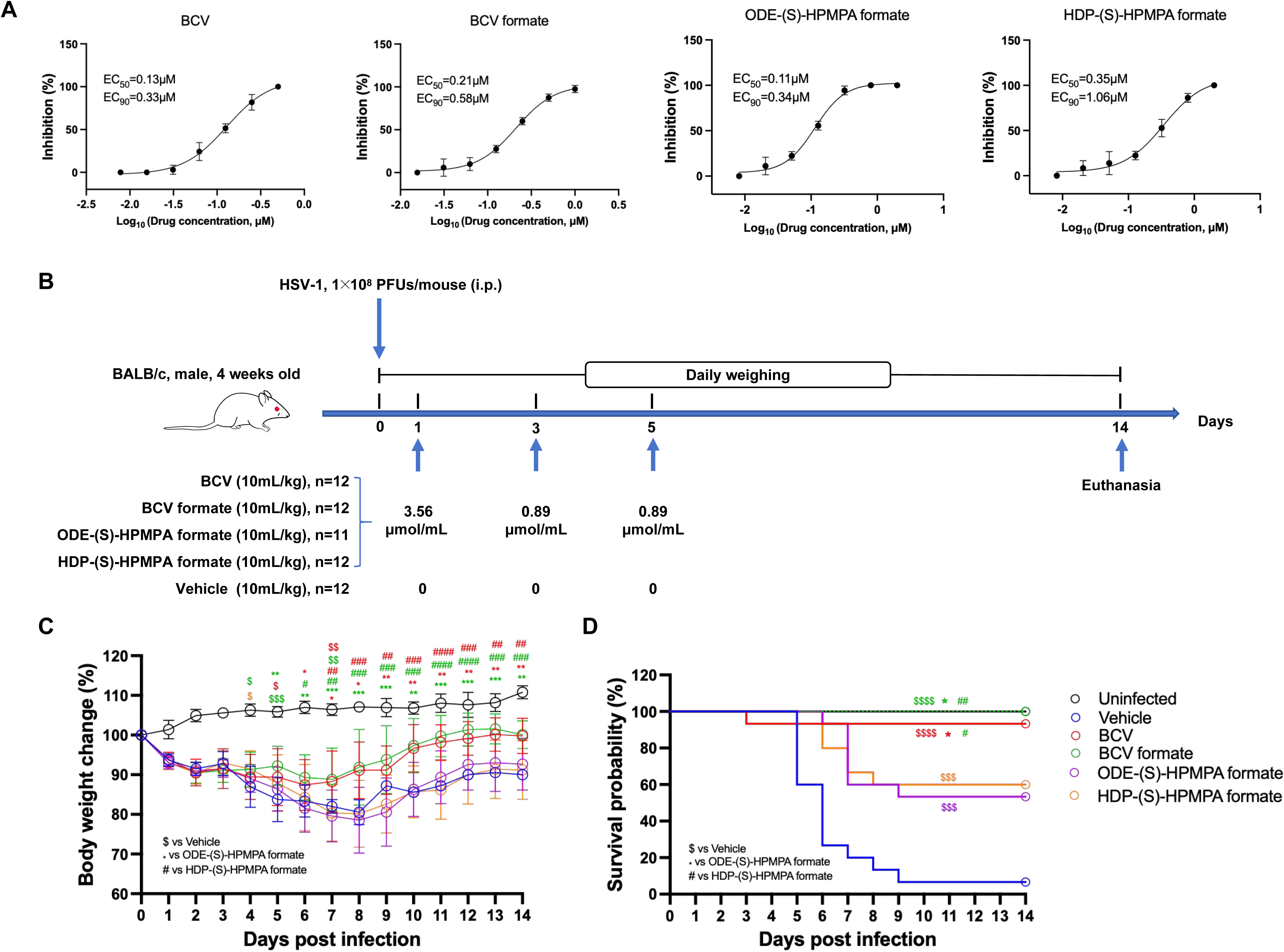
Evaluations of in vivo and in vitro anti-HSV-1 potencies of BCV and the candidate prodrugs. (**A**) Vero cells were infected with HSV-1 stain 17 and treated with indicated concentrations of BCV or the candidate compounds. All compounds were tested in triplicated wells at each concentration. The solvent was used as the non-treated control. The antiviral effects were determined by calculating the reduction in plaque formation. Data are presented as mean ± SD. (**B**) The schematic illustration of in vivo experiment design. (**C-D**) Weight loss and survival rates of mice after being intra peritoneally infected with HSV-1. Mice received the solvent served as the non-treated control. The data of weight loss are presented as mean ± SD. Weight loss ≥30% of initial body weight was recorded as ethical death and was therefore not represented in the figure. Statistical significance is indicated as follows: *, $, #: p < 0.05; **, $$, ##: p < 0.01; ***, $$$, ###: p < 0.001; ****, $$$$, ####: p < 0.0001.

Inspired by this observation, we were curious whether a combined regimen of purine and pyrimidine analogs would provide more balanced protection against orthopoxvirus and herpesvirus. To test this hypothesis, we treated vaccinia infected mice with the equal molar mixture of BCV formate and ODE-(S)-HPMPA formate, and treated HSV-1 infected mice with the equal molar mixture of BCV formate and ODE-(S)-HPMPA formate (Supplementary figure 6A). The total dosage for each mixed regimen was maintained in consistency with the aforementioned single-drug regimen. The result showed that all mice survived after being treated with the combined regimens (Supplementary figure 6B), which was obviously better than any single-drug treatment (Figure 4 and Figure 6).

### Three candidate compounds exhibited better in vitro anti-hAdV activity than BCV

BCV has been used compassionately to treat adenovirus infection for pediatric patients receiving hematopoietic stem cell transplantation [42]. In this study, we compared the anti-hAdV potencies between three candidate prodrugs and BCV using clinically isolated strains of hAdV type C2 and type B21. Generally, the EC_50_ values of the three candidates, especially those of BCV formate and ODE-(S)-HPMPA formate, were much lower than the EC_50_ values BCV (Figure 7). Considering that the liver toxicity of ODE-(S)-HPMPA formate were significantly lower than BCV formate (Supplementary figure 4), we thought ODE-(S)-HPMPA formate could also be good candidate therapeutic for hAdV infections.

**Figure 7.**
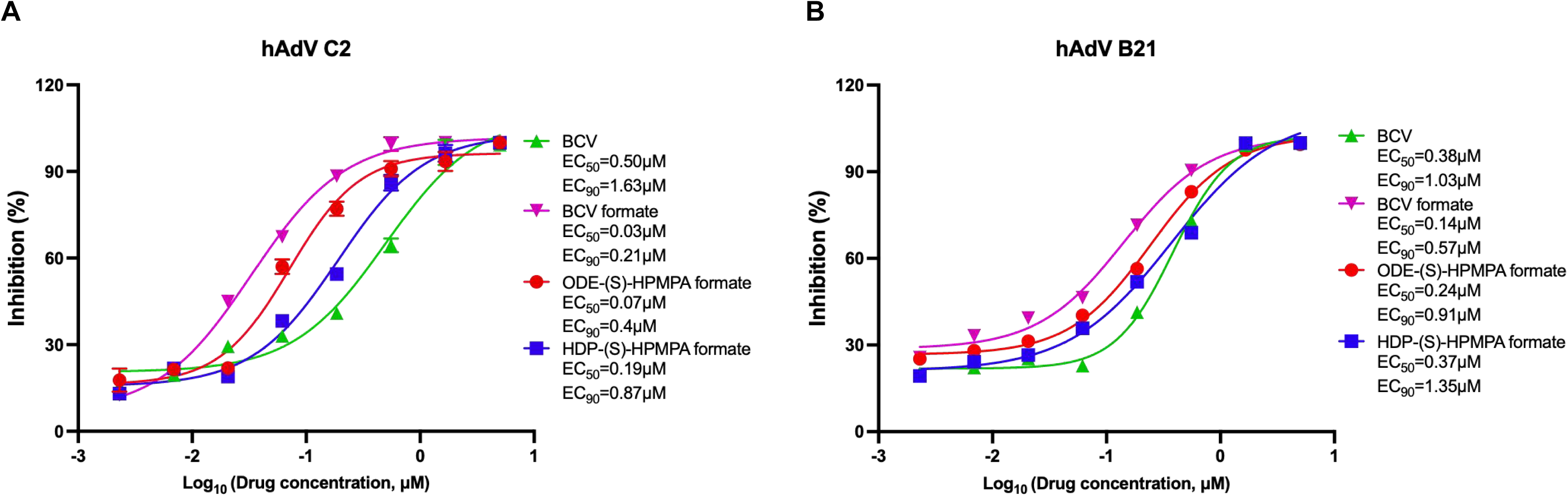
In vitro evaluation of anti-hAdV activities of BCV and the candidate compounds. Vero cells were infected with either hAdV C2 (**A**) or hAdV B21 (**B**) and subsequently treated with indicated concentrations of BCV or the candidate prodrugs. All compounds were tested in quadruplicate at each concentration. The solvent was used as the non-treated control. Data are presented as mean ± SD.

## Discussion

Nucleoside/nucleotide analogues (NAs) have been widely used to treat cancer and viral diseases [43]. Given that the catalytic units of viral polymerases are conserved in their structures [44], NAs are thought to be an important class of broad-spectrum antiviral agents [41]. Since the first anti-viral NA was approved in 1969, more than two dozen NAs have been licensed to treat viral infections [45]. However, only limited number of viral diseases are targeted [3]. Among the six families of double stranded DNA viruses (Hepadnaviridae, Polyomaviridae, Papillomaviridae, Adenoviridae, Herpesviridae and Poxviridae) known to cause human diseases [46], only five viruses from three families (Hepadnaviridae, Herpesviridae and Poxviridae) can be treated with approved NAs [3, 4]. BCV is currently the only NA that can be used to treat human Mpox disease under an FDA-authorized single-patient e-IND, while its efficacy is still inconclusive [47]. Bottlenecks, such as toxicity, poor oral bioavailability and limited translatability, restrain NAs from being applied to more viral diseases [41, 45]. In this work, we and collaborators synthesized a new precursor of CDV and two novel precursors of (S)- HPMPA and evaluated their inhibitory activities against multiple double-stranded DNA viruses, including orthopoxviruses, HSV-1 (strain 17) and human adenoviruses.

Through the experiments shown here, we found that BCV and three candidate prodrugs exhibited varied antiviral potencies against different DNA viruses. The anti-orthopoxvirus potency of BCV formate was non-inferior to BCV, by contrast the two prodrugs of (S)-HPMPA were remarkably more effective than BCV in suppressing orthopoxviruses both in vitro and in vivo. ODE-(S)-HPMPA formate was the best-performing anti-orthopoxvirus compound, which displayed the most robust anti-orthopoxvirus activity and the lowest hepatotoxicity. ODE-(S)-HPMPA formate and HDP-(S)-HPMPA formate were designed based on previously reported ODE-(S)- HPMPA and HDP-(S)-HPMPA [30, 33]. We did not test the latter compounds in parallel in this study, but compared the data to that available in the literature [33], both ODE-(S)- HPMPA formate and HDP-(S)-HPMPA formate are much more efficacious than their counterparts in protecting mice from lethal vaccinia infection (Supplementary table 3). The fact that ODE-(S)-HPMPA formate was more efficacious than HDP-(S)-HPMPA formate in protecting mice against vaccinia infection might partly because ODE-(S)- HPMPA formate distributed to lung more efficiently than HDP-(S)-HPMPA formate (Supplementary figure 3).

A second key finding of this work is that BCV and BCV formate provided significantly stronger protection against lethal HSV-1 infection than ODE-(S)-HPMPA formate and HDP-(S)-HPMPA formate in mice. The primary anti-viral mechanism of NAs is that they resemble natural nucleos(t)ides and act by being incorporated into the nascent DNA chain [48, 49], however the mechanism underlying the varied protective effects against orthopoxvirus and HSV-1 is not clear. As herpesvirus genome contains obviously more GC than orthopoxvirus genome, we speculate that the cytidine analogs (BCV and BCV formate) might be more effective in suppressing herpesviruses, while adenine analogs, ODE-(S)-HPMPA formate and HDP-(S)-HPMPA formate, might inhibit orthopoxviruses more efficiently. Consistent with this hypothesis, we proved that combined regimens of cytidine and adenine analogs could provide excellent protection against both vaccinia and HSV-1 infection in mice, indicating that combination of purine and pyrimidine analogs might be a rational and promising strategy to develop a broad-spectrum antiviral therapy.

Finally, we tested the anti-hAdV potencies of the candidate prodrugs through in vitro experiments. Adenoviruses can cause life-threatening disease not only in immunocompromised patients but also in the healthy, although the latter occurs less frequently [50, 51]. Both CDV and BCV have been compassionately used to treat hAdV infections in children receiving hematopoietic cell transplantation [42, 52], but there are no officially approved therapeutics. Here we showed ODE-(S)-HPMPA formate could be a good anti-hAdV agent, which exhibited a good safety profile and relatively stronger inhibitory effect than BCV.

A major limitation of this work is that we did not test the in vivo anti-MPXV and anti-hAdV effects of the candidate prodrugs, primarily because no appropriate animal models are available to us. Evaluations using an MPXV infected rhesus macaque model [53] and an hAdV infected human lung organoid model [54] may provide more relevant data before moving any of the candidate prodrugs into clinical trials.

## Author contributions

Yanmin Wan, Wenhong Zhang and Youchun Wang jointly supervised the study. Yanmin Wan, Zhaoqin Zhu and Chao Qiu designed the experiments. Yanmin Wan, Yifan Zhang, Cuiyuan Guo, Jiasheng Lu, Yanan Zhou, Jiaojiao Zheng, Fahui Dai, Xiaoyang Cheng and Kunlu Deng conducted the experiments. Yanmin Wan, Yifan Zhang and Cuiyuan Guo analyzed the data and drafted the manuscript. Yanmin Wan, Jiasheng Lu, Zhaoqin Zhu and Wanhai Wang revised the manuscript. All authors approved the submitted version.

## Acknowledgments

We thank Quanlin Xue from Fudan University for kindly sharing HSV-1 (strain 17) with us. This work was supported in part by the National Natural Science Foundation of China (Grant No. 32270986), the major project of Study on Pathogenesis and Epidemic Prevention Technology System by the Ministry of Science and Technology of China (Grant No. 2021YFC2302500) and the National Key Research and Development Program of China (Grant No. 2022YFC2009802).

## Supplementary figure legends

**Supplementary figure 1.**
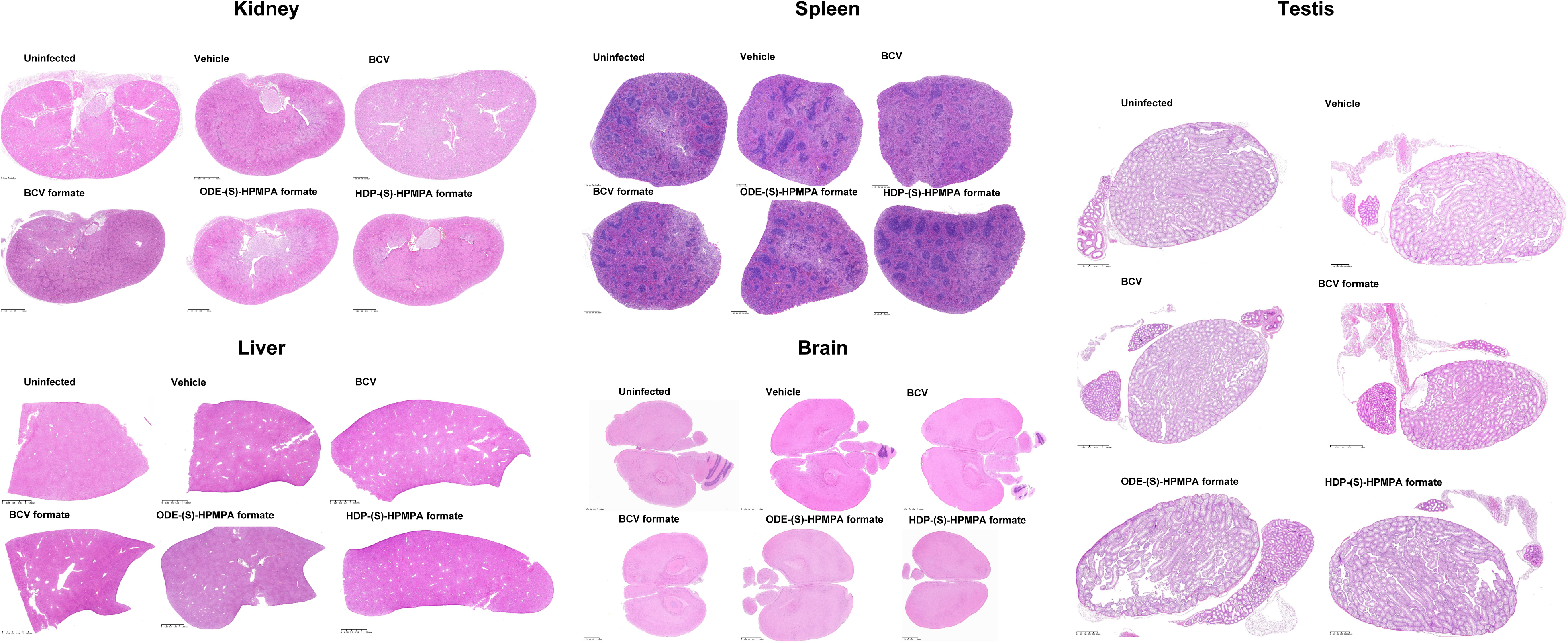
No signs of infection-caused or drug-related pathology were detected in key organs post Tiantan vaccinia infection. Histopathological examination of the kidneys, spleen, testis, liver, and brain in mice at 7 days post Tiantan vaccinia infection.

**Supplementary figure 2.**
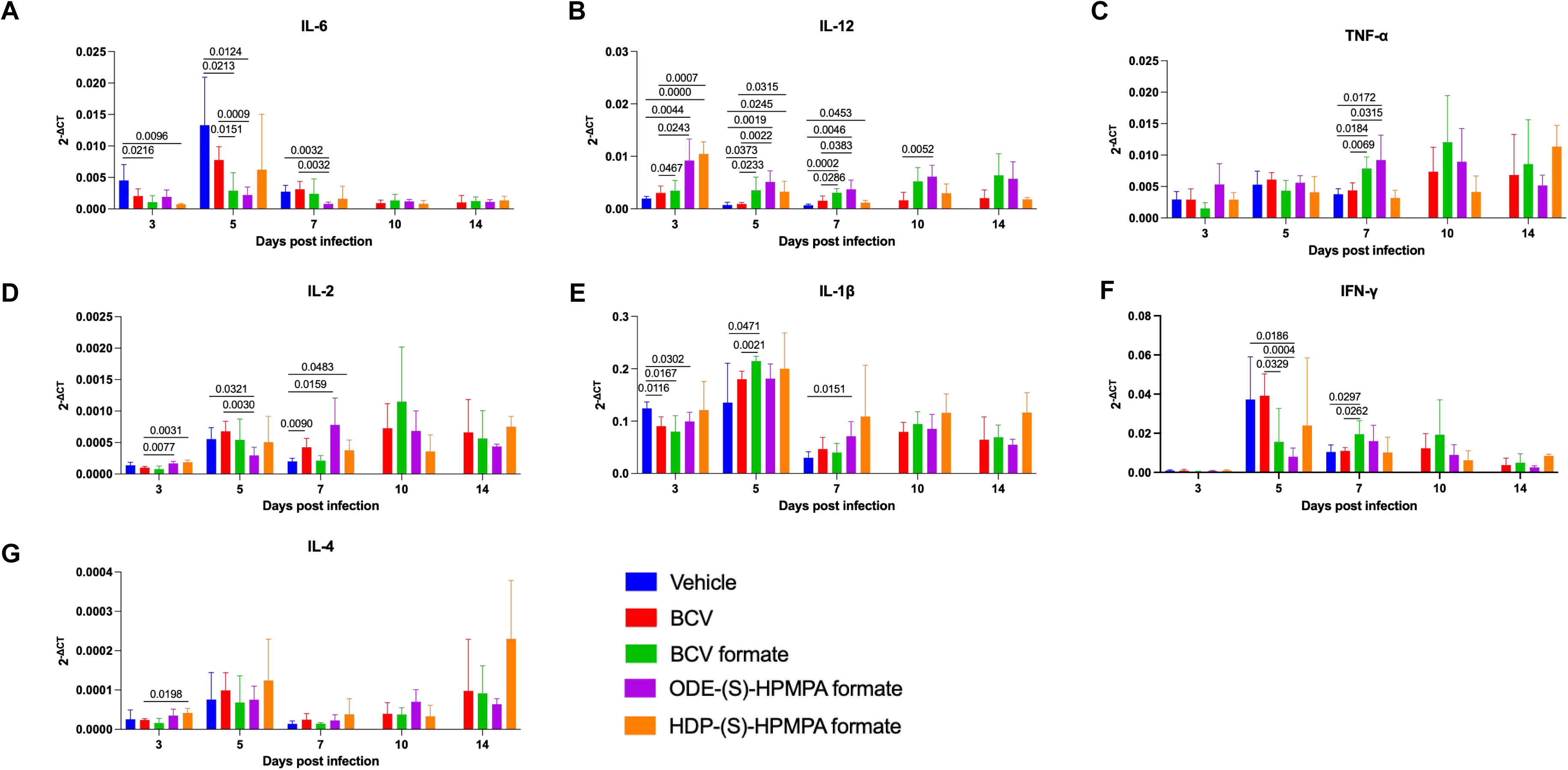
Drug treatment impacted pulmonary cytokine responses in mice following intranasal infection with Tiantan vaccinia. The transcriptional levels of IL-6 (**A**), IL-12 (**B**), TNF-α (**C**), IL-2 (**D**), IL-1β (**E**), IFN-γ (**F**), IL-4 (**G**) in lung tissues were assessed at 3, 5, 7, 10, and 14 days post-infection with the Tiantan vaccinia virus.

**Supplementary figure 3.**
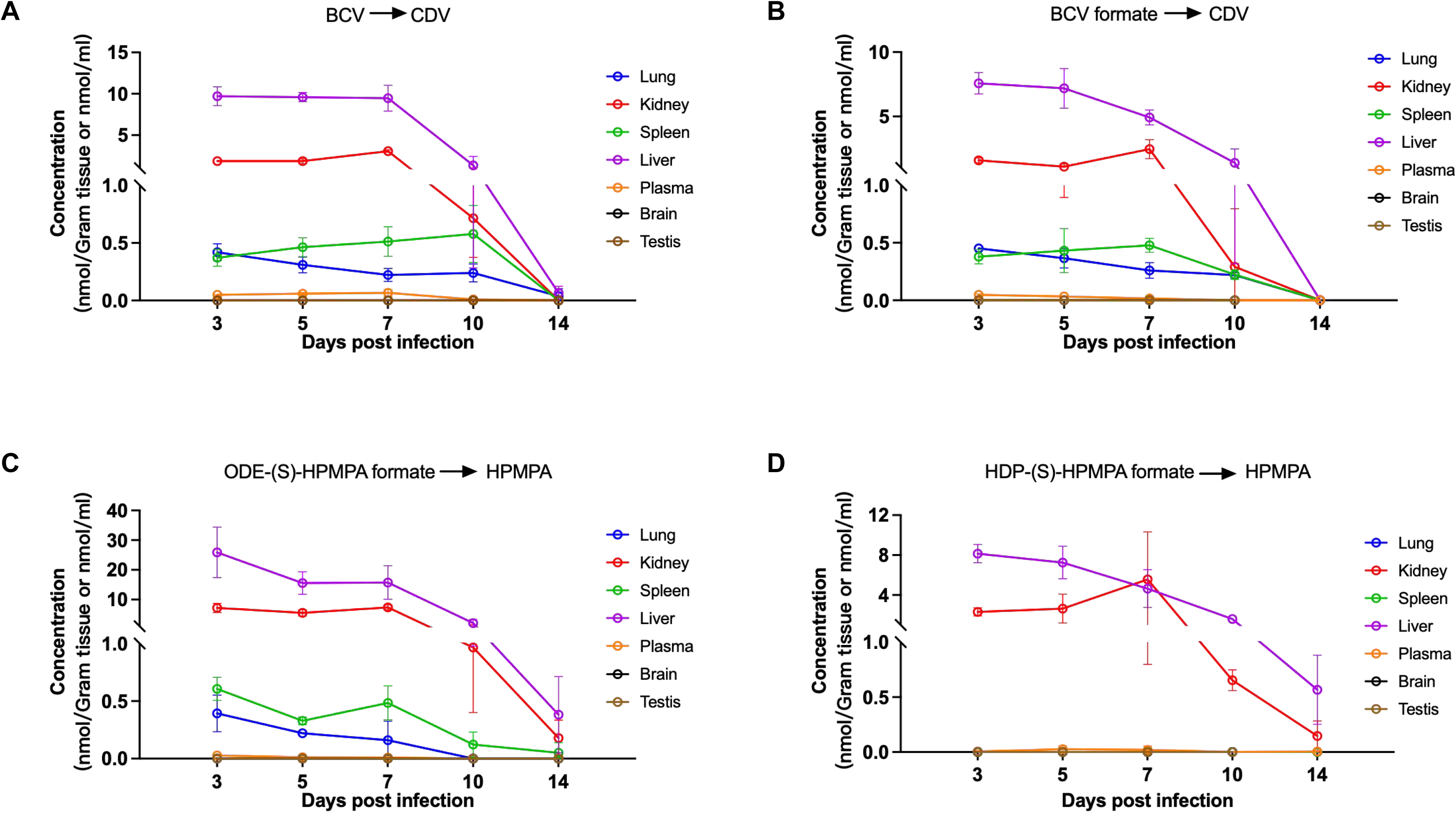
Monitoring in vivo drug metabolites throughout the treatment course. (**A-D**) The concentrations of drug metabolites in mouse plasma, lung, kidney, spleen and liver were measured using HPLC-MS at 3, 5, 7, 10 and 14 days post infection, respectively (3 mice per group). Metabolites of all drugs were non-detectable in the brain and testis at any time point. Data are presented as mean ± SD.

**Supplementary figure 4.**
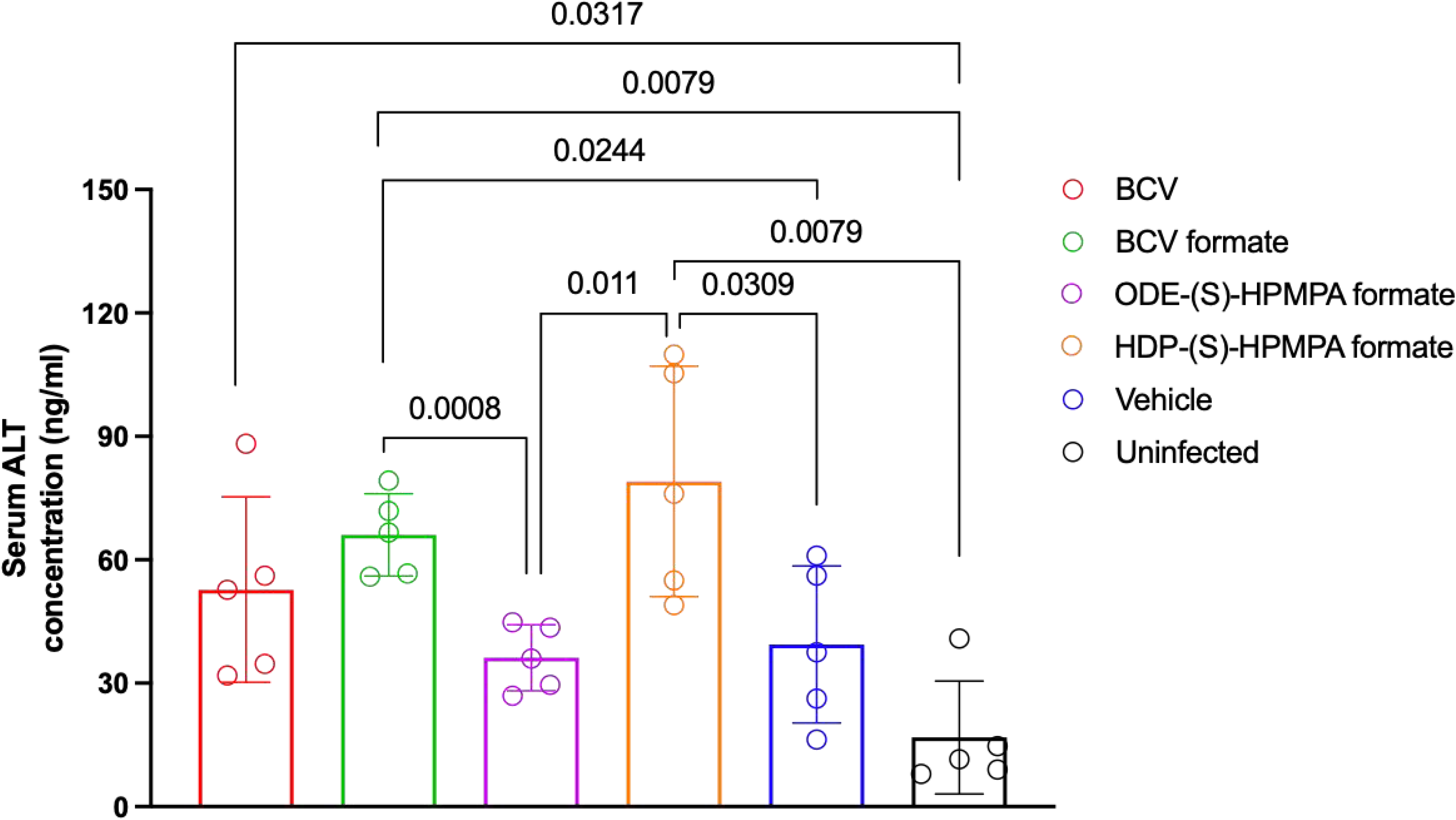
Detection of ALT levels in mouse serum. Serum ALT levels were measured at 2 days after the third dose drug treatment (n=5). Data are presented as mean ± SD.

**Supplementary figure 5.**
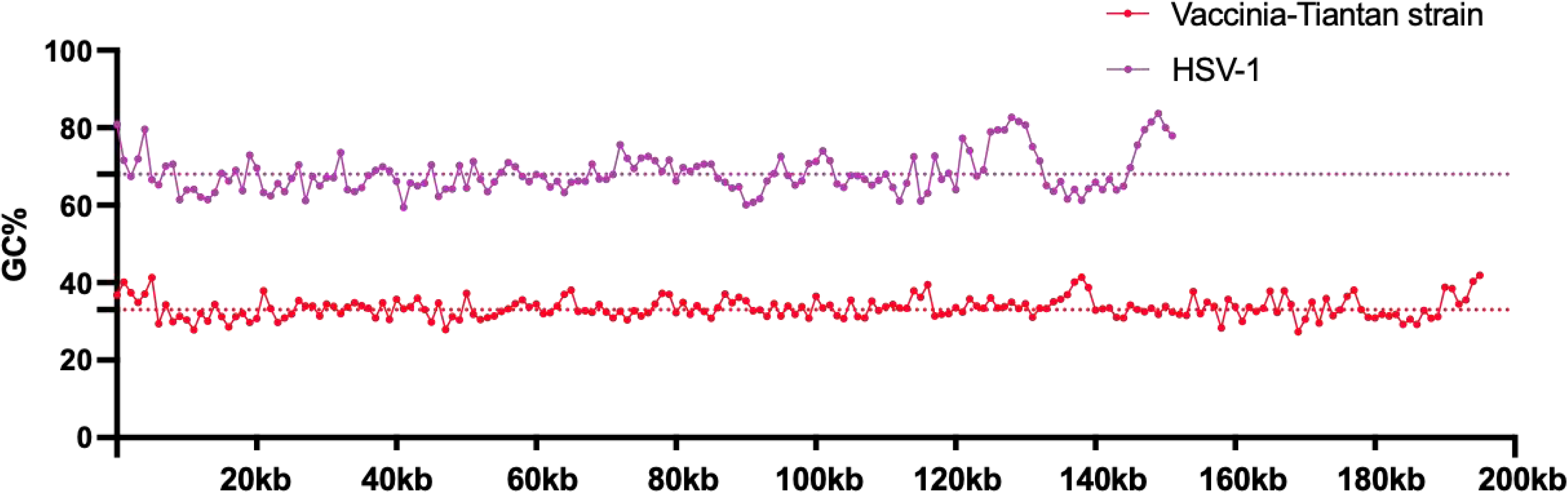
Comparison of genome GC content between Tiantan vaccinia and HSV-1. A representative Tiantan vaccinia genome sequence (accession number: KC207811.1) and a representative HSV-1 genome sequence (accession number: X14112.1) were retrieved from GenBank. The GC content was analyzed using a sliding window approach in the Biostrings package of R package, with a window size of 1000 base pairs.

**Supplementary figure 6.**
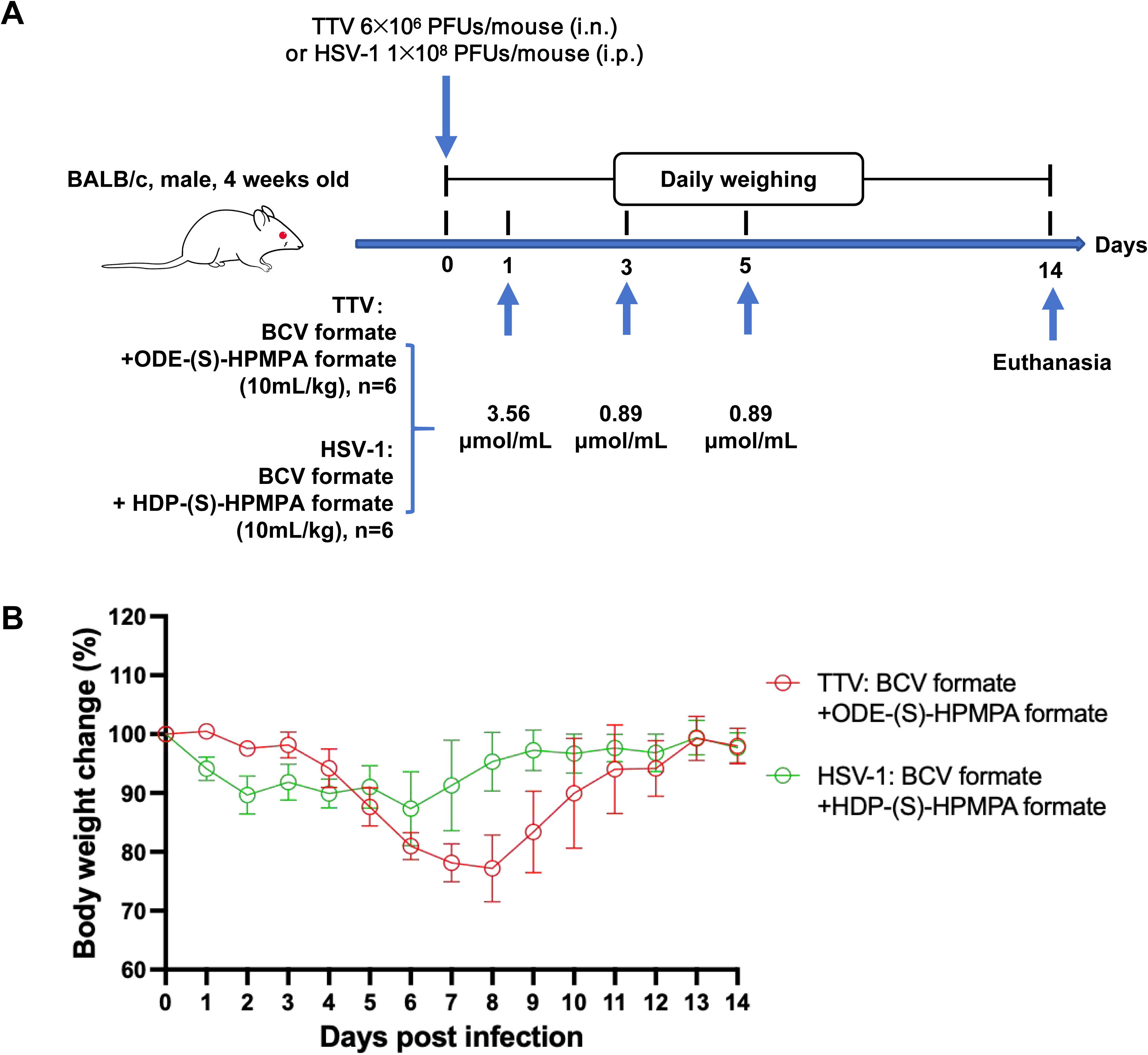
Adenine and cytidine analog combinations showed complete protection against both vaccinia and HSV-1 infections. (**A**) The schematic illustration of experimental design. (**B**) Weight loss of mice after being infected with Tiantan vaccinia or HSV-1. Data are presented as mean ± SD.

**Supplementary table 1.**
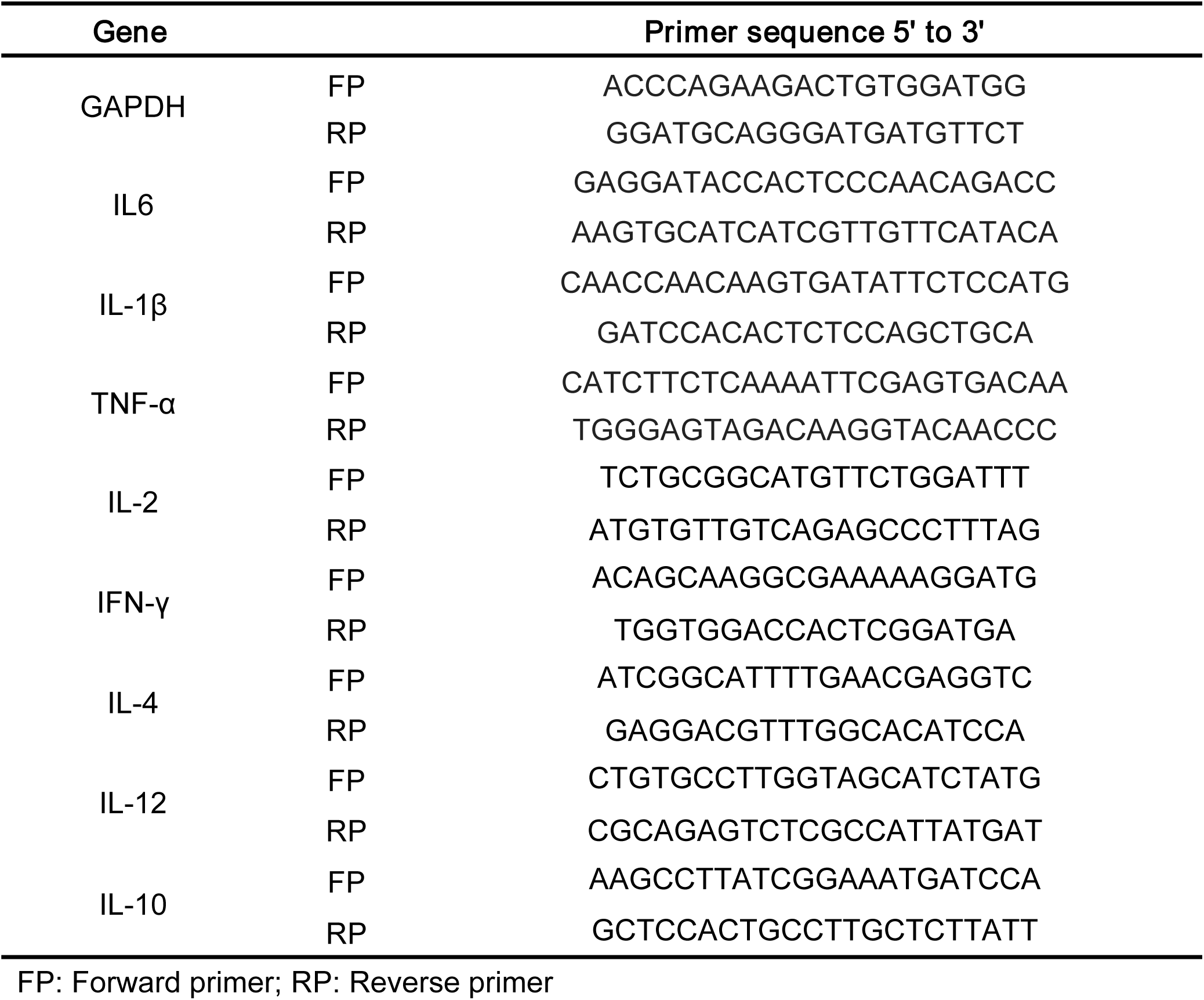
Primer sequences used for quantitative PCR assays of murine cytokines.

**Supplementary table 2.** GC content analyses of representative genomes of orthopoxviruses and herpesviruses.

| Virus | GenBank Number | GC% |
| --- | --- | --- |
| Vaccinia virus strain TianTan | KC207811.1 | 33 |
| Vaccinia virus WR | AY243312.1 | 33 |
| Monkeypox virus clade IIb B.1 | ON676708.1 | 33 |
| Monkeypox virus clade Ib | PP601207.1 | 33 |
| Monkeypox virus clade Ia | PP601206.1 | 33 |
| Cowpox virus | NC_003663.2 | 33 |
| Variola virus | PP405600.1 | 33 |
| Human herpesvirus 1 | X14112.1 | 68 |
| Human herpesvirus 2 | NC_001798.2 | 70 |
| Human cytomegalovirus | X17403.1 | 57 |
| Epstein-Barr virus | V01555.2 | 60 |

**Supplementary table 3.**
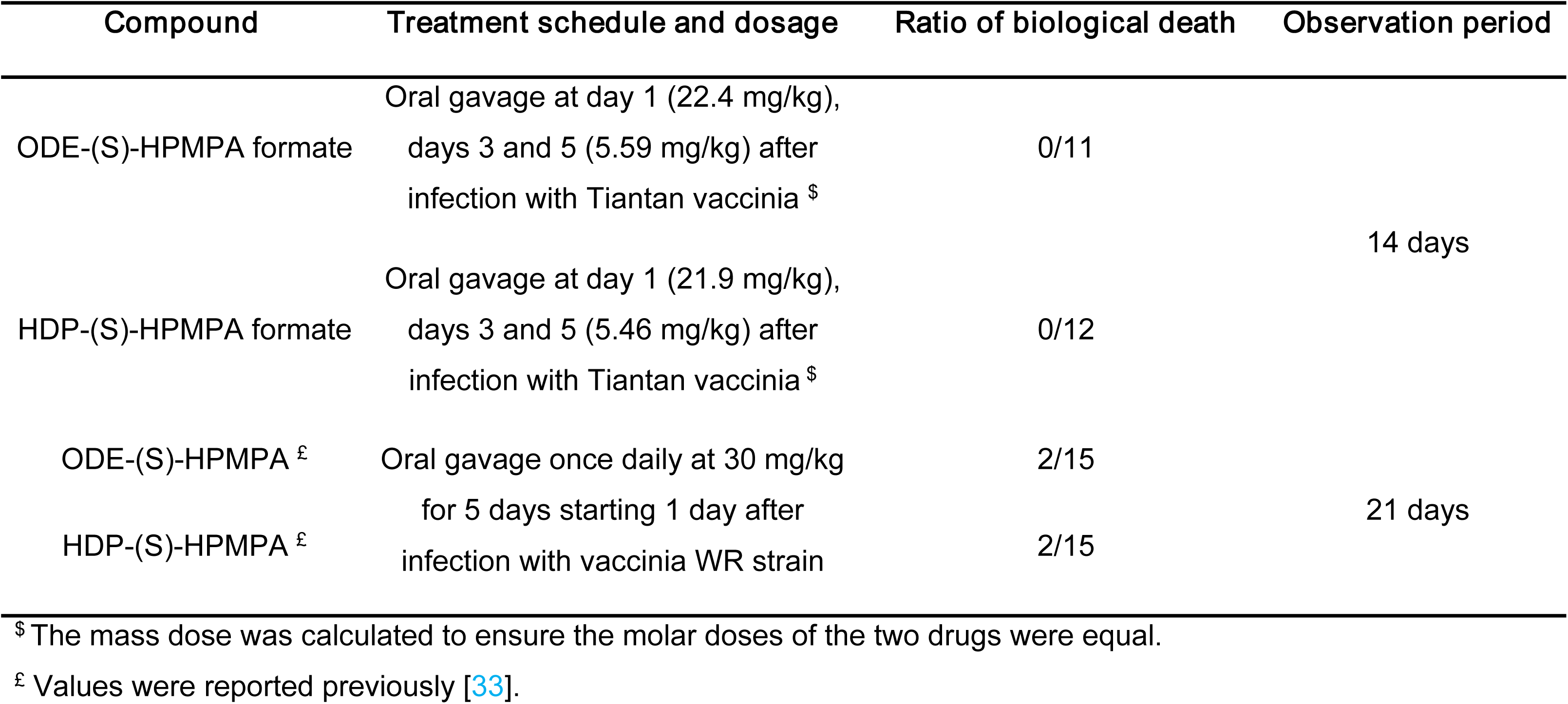
Both ODE-(S)-HPMPA formate and HDP-(S)-HPMPA formate are more efficacious than ODE- (S)-HPMPA and HDP-(S)-HPMPA in protecting mice from lethal vaccinia infection.

## References

1. Lu, L., et al., Antivirals with common targets against highly pathogenic viruses. Cell, 2021. 184(6): p. 1604–1620.

2. Peng, S., et al., Progression of Antiviral Agents Targeting Viral Polymerases. Molecules, 2022. 27(21).

3. Palazzotti, D., et al., Small Molecule Drugs Targeting Viral Polymerases. Pharmaceuticals (Basel), 2024. 17(5).

4. Chan-Tack, K., et al., Benefit-risk assessment for brincidofovir for the treatment of smallpox: U.S. Food and Drug Administration’s Evaluation. Antiviral Res, 2021. 195: p. 105182.

5. Magee, W.C. and D.H. Evans, The antiviral activity and mechanism of action of (S)-[3-hydroxy-2-(phosphonomethoxy)propyl] (HPMP) nucleosides. Antiviral Res, 2012. 96(2): p. 169–80.

6. Bojkova, D., et al., Drug Sensitivity of Currently Circulating Mpox Viruses. N Engl J Med, 2023. 388(3): p. 279–281.

7. Frenois-Veyrat, G., et al., Tecovirimat is effective against human monkeypox virus in vitro at nanomolar concentrations. Nat Microbiol, 2022. 7(12): p. 1951–1955.

8. Byrareddy, S.N., et al., Potential therapeutic targets for Mpox: the evidence to date. Expert Opin Ther Targets, 2023. 27(6): p. 419–431.

9. Duraffour, S., G. Andrei, and R. Snoeck, Tecovirimat, a p37 envelope protein inhibitor for the treatment of smallpox infection. IDrugs, 2010. 13(3): p. 181–91.

10. Shamim, M.A., et al., The use of antivirals in the treatment of human monkeypox outbreaks: a systematic review. Int J Infect Dis, 2023. 127: p. 150–161.

11. Adler, H., et al., Clinical features and management of human monkeypox: a retrospective observational study in the UK. Lancet Infect Dis, 2022. 22(8): p. 1153–1162.

12. Barnes, A.H., et al., Mpox: Special Considerations in the Immunocompromised Host. Current Treatment Options in Infectious Diseases, 2022. 14(4): p. 43–66.

13. Imran, M., et al., Oral Brincidofovir Therapy for Monkeypox Outbreak: A Focused Review on the Therapeutic Potential, Clinical Studies, Patent Literature, and Prospects. Biomedicines, 2023. 11(2).

14. Grosenbach, D.W., et al., Oral Tecovirimat for the Treatment of Smallpox. N Engl J Med, 2018. 379(1): p. 44–53.

15. Duraffour, S., et al., ST-246 is a key antiviral to inhibit the viral F13L phospholipase, one of the essential proteins for orthopoxvirus wrapping. J Antimicrob Chemother, 2015. 70(5): p. 1367–80.

16. Smith, T.G., et al., Tecovirimat Resistance in Mpox Patients, United States, 2022-2023. Emerg Infect Dis, 2023. 29(12): p. 2426–2432.

17. Alarcón, J., et al., An Mpox-Related Death in the United States. N Engl J Med, 2023. 388(13): p. 1246–1247.

18. Lenharo, M., Hopes dashed for drug aimed at monkeypox virus spreading in Africa. Nature, 2024. 632(8027): p. 965.

19. Andrei, G. and R. Snoeck, Cidofovir Activity against Poxvirus Infections. Viruses, 2010. 2(12): p. 2803–30.

20. Lurain, N.S. and S. Chou, Antiviral drug resistance of human cytomegalovirus. Clin Microbiol Rev, 2010. 23(4): p. 689–712.

21. Farlow, J., et al., Comparative whole genome sequence analysis of wild-type and cidofovir-resistant monkeypoxvirus. Virol J, 2010. 7: p. 110.

22. Silva, N.I.O., et al., Here, There, and Everywhere: The Wide Host Range and Geographic Distribution of Zoonotic Orthopoxviruses. Viruses, 2020. 13(1).

23. Zhang, Y., et al., Potential threat of human pathogenic orthopoxviruses to public health and control strategies. J Biosaf Biosecur, 2023. 5(1): p. 1–7.

24. Shchelkunov, S.N., An increasing danger of zoonotic orthopoxvirus infections. PLoS Pathog, 2013. 9(12): p. e1003756.

25. Shchelkunova, G.A. and S.N. Shchelkunov, Smallpox, Monkeypox and Other Human Orthopoxvirus Infections. Viruses, 2022. 15(1).

26. De Clercq, E. and A. Holý, Acyclic nucleoside phosphonates: a key class of antiviral drugs. Nat Rev Drug Discov, 2005. 4(11): p. 928–40.

27. Tollefson, A.E., et al., Oral USC-093, a novel homoserinamide analogue of the tyrosinamide (S)-HPMPA prodrug USC-087 has decreased nephrotoxicity while maintaining antiviral efficacy against human adenovirus infection of Syrian hamsters. Antiviral Res, 2024. 222: p. 105799.

28. Luo, M., et al., Amidate Prodrugs of Cyclic 9-(S)-[3-Hydroxy-2-(phosphonomethoxy)propyl]adenine with Potent Anti-Herpesvirus Activity. ACS Med Chem Lett, 2018. 9(4): p. 381–385.

29. Beadle, J.R., Synthesis of cidofovir and (S)-HPMPA ether lipid prodrugs. Curr Protoc Nucleic Acid Chem, 2007. Chapter 15: p. Unit 15.2.

30. Quenelle, D.C., et al., Effect of oral treatment with (S)-HPMPA, HDP-(S)-HPMPA or ODE-(S)-HPMPA on replication of murine cytomegalovirus (MCMV) or human cytomegalovirus (HCMV) in animal models. Antiviral Res, 2008. 79(2): p. 133–5.

31. Dal Pozzo, F., et al., In vitro evaluation of the anti-orf virus activity of alkoxyalkyl esters of CDV, cCDV and (S)-HPMPA. Antiviral Res, 2007. 75(1): p. 52–7.

32. Morrey, J.D., et al., Alkoxyalkyl esters of 9-(s)-(3-hydroxy-2-phosphonomethoxypropyl) adenine are potent and selective inhibitors of hepatitis B virus (HBV) replication in vitro and in HBV transgenic mice in vivo. Antimicrob Agents Chemother, 2009. 53(7): p. 2865–70.

33. Quenelle, D.C., et al., Effect of oral treatment with hexadecyloxypropyl-[(S)-9-(3-hydroxy-2-phosphonylmethoxypropyl)adenine] [(S)-HPMPA] or octadecyloxyethyl-(S)- HPMPA on cowpox or vaccinia virus infections in mice. Antimicrob Agents Chemother, 2007. 51(11): p. 3940–7.

34. Beadle, J.R., et al., Synthesis and antiviral evaluation of alkoxyalkyl derivatives of 9-(S)- (3-hydroxy-2-phosphonomethoxypropyl)adenine against cytomegalovirus and orthopoxviruses. J Med Chem, 2006. 49(6): p. 2010–5.

35. Ruiz, J., et al., Synthesis, metabolic stability and antiviral evaluation of various alkoxyalkyl esters of cidofovir and 9-(S)-[3-hydroxy-2- (phosphonomethoxy)propyl]adenine. Bioorg Med Chem, 2011. 19(9): p. 2950–8.

36. Grosche, L., et al., Herpes Simplex Virus Type 1 Propagation, Titration and Single-step Growth Curves. Bio Protoc, 2019. 9(23): p. e3441.

37. Hartline, C.B., et al., A standardized approach to the evaluation of antivirals against DNA viruses: Orthopox-, adeno-, and herpesviruses. Antiviral Res, 2018. 159: p. 104–112.

38. Painter, W., et al., First pharmacokinetic and safety study in humans of the novel lipid antiviral conjugate CMX001, a broad-spectrum oral drug active against double-stranded DNA viruses. Antimicrob Agents Chemother, 2012. 56(5): p. 2726–34.

39. Wang, X., et al., Immune Correlates of Disseminated BCG Infection in IL12RB1-Deficient Mice. Vaccines (Basel), 2022. 10(7).

40. Potmesil, P., et al., Nucleotide analogues with immunobiological properties: 9-[2-Hydroxy-3-(phosphonomethoxy)propyl]-adenine (HPMPA), -2,6-diaminopurine (HPMPDAP), and their N6-substituted derivatives. Eur J Pharmacol, 2006. 540(1-3): p. 191–9.

41. Karim, M., C.W. Lo, and S. Einav, Preparing for the next viral threat with broad-spectrum antivirals. J Clin Invest, 2023. 133(11).

42. Perruccio, K., et al., Safety and efficacy of brincidofovir for Adenovirus infection in children receiving allogeneic stem cell transplantation: an AIEOP retrospective analyses. Bone Marrow Transplant, 2021. 56(12): p. 3104–3107.

43. Jordheim, L.P., et al., Advances in the development of nucleoside and nucleotide analogues for cancer and viral diseases. Nature Reviews Drug Discovery, 2013. 12(6): p. 447–464.

44. Peersen, O.B., A Comprehensive Superposition of Viral Polymerase Structures. Viruses, 2019. 11(8).

45. Jordheim, L.P., et al., Advances in the development of nucleoside and nucleotide analogues for cancer and viral diseases. Nat Rev Drug Discov, 2013. 12(6): p. 447–64.

46. Modrow, S., et al., Viruses with a Double-Stranded DNA Genome, in Molecular Virology, S. Modrow, et al., Editors. 2013, Springer Berlin Heidelberg: Berlin, Heidelberg. p. 625–873.

47. Li, P., et al., Clinical Features, Antiviral Treatment, and Patient Outcomes: A Systematic Review and Comparative Analysis of the Previous and the 2022 Mpox Outbreaks. J Infect Dis, 2023. 228(4): p. 391–401.

48. Nucleoside Analogues, in LiverTox: Clinical and Research Information on Drug-Induced Liver Injury. 2012, National Institute of Diabetes and Digestive and Kidney Diseases: Bethesda (MD).

49. Kamzeeva, P.N., et al., Recent Advances in Molecular Mechanisms of Nucleoside Antivirals. Curr Issues Mol Biol, 2023. 45(8): p. 6851–6879.

50. Khanal, S., P. Ghimire, and A.S. Dhamoon, The Repertoire of Adenovirus in Human Disease: The Innocuous to the Deadly. Biomedicines, 2018. 6(1).

51. Dotan, M., et al., Adenovirus can be a serious, life-threatening disease, even in previously healthy children. Acta Paediatr, 2022. 111(3): p. 614–619.

52. Hiwarkar, P., et al., Brincidofovir is highly efficacious in controlling adenoviremia in pediatric recipients of hematopoietic cell transplant. Blood, 2017. 129(14): p. 2033–2037.

53. Aid, M., et al., Mpox infection protects against re-challenge in rhesus macaques. Cell, 2023. 186(21): p. 4652–4661.e13.

54. Zhao, S., et al., Generation of Human Embryonic Stem Cell-Derived Lung Organoids for Modeling Infection and Replication Differences between Human Adenovirus Types 3 and 55 and Evaluating Potential Antiviral Drugs. J Virol, 2023. 97(5): p. e0020923.

